# Astrocyte diversity and aging in the mouse lemur primate brain

**DOI:** 10.64898/2026.01.26.701678

**Authors:** Lolie Garcia, Léo Dupuis, Fanny Petit, Suzanne Lam, Jean-Luc Picq, Marc Dhenain

## Abstract

Astrocytes play key roles in maintaining brain homeostasis, metabolism, and neurovascular integrity, yet their diversity and age-related modulation remain insufficiently understood, particularly across primate lineages. While rodent studies have generated extensive knowledge, notable species differences highlight the need for comparative analyses in non-human primates. The gray mouse lemur (*Microcebus murinus*), a small primate widely used in aging research, offers a valuable but underexplored model for studying astroglial aging. In this study, we characterized astrocyte distribution, morphology, and reactivity in 17 mouse lemurs aged 1.0–11.5 years using GFAP and vimentin immunohistochemistry. We identified marked regional and morphological heterogeneity, with dense astrocytic labeling in white matter, hippocampus, and sparse but diverse cortical populations. Distinct astrocyte subtypes—including fibrous, protoplasmic, projection, pial and subpial interlaminar, radial glia-like cells, tanycytes—were documented. Varicosity-bearing processes were common across multiple astroglial subtypes and may indicate altered physiological states. Quantitative analyses revealed pronounced age-related increases in astrocytic reactivity, particularly in white matter and interlaminar astrocytes. Cortical and hippocampal changes were comparatively modest. These findings indicate region-specific astrocytic vulnerability during aging and support the translational value of the mouse lemur for investigating glial aging in primates.

**Main Points:**

- The mouse lemur is the smallest primate on earth with a key role to understand primate brain characteristics.
- We characterized seven different astrocyte subtypes: from fibrous to primate-specific astrocyte as interlaminar astrocytes in different brain regions of this primate.
- Varicosities were reported in different astrocyte subtypes found close to brain borders.
- Main age-related changes concerned fibrous astrocytes in the white matter and interlaminar astrocytes at the cortical border.

## 1. Introduction

Astrocytes play diverse and essential roles to regulate the extracellular brain environment and support brain functions, including maintaining ion and water homeostasis, preserving the blood– brain barrier, regulating cerebral blood flow, managing neurotransmitter, synaptic, and metabolic regulations ^1-7^. They create a syncitium or astroglia network that is critical for organizing brain function including through calcium waves ^8-10^.

In the mammalian central nervous system, various astrocyte subtypes presenting with different morphology, physiology and metabolism have long been recognized ^11-14^. Different subtypes can populate a given brain region or different brain regions. The earliest identified astrocyte types are protoplasmic and fibrous astrocytes ^15^. Protoplasmic astrocytes, the most abundant astrocyte population in gray matter, are primarily located in cerebral cortex and the hippocampus. Their highly branched cell bodies that wrap and envelop blood vessels and enable contacts with numerous synapses, support a neuromodulatory function ^12^. In contrast, fibrous astrocytes are aligned with white matter tracts. They are smaller, less complex in their branching, and can interact with nodes of Ranvier ^16^. Other astrocytes are also described as polarized gray matter astrocytes, reported for example in the dentate gyrus of rodents and other species, where they extend most of their processes towards the hippocampal fissure ^17^. In addition to this regional variability, animal species differences have been reported with some astrocyte subtypes reported only in some species as primates or carnivores ^13,14,18^. A typical example is interlaminar astrocytes (ILA). These astrocytes that are reported only in primates and carnivores have a cell body located in the upper cortical layer 1 and send long, frequently unbranched processes towards deeper layers, terminating in either layer 3 or 4 ^13,18^. Comparative studies across primate species indicate that their morphological features vary in different primates with increased morphological complexity reported in anthropoid primates compared to strepsirrhines ^19^. Varicose projection astrocytes (VPAs) are another category of astrocytes that show differences between species. They are characterized by 1–5 very long fibers, covered with regularly spaced varicosities, that extend in multiple directions across the cerebral cortex ^12,13,20^. They reside in the deep cortical layers, close to the white matter, and are also referred to as polarized astrocytes ^19^. Although VPAs were originally considered a primate-specific subtype of astrocytes, recent work suggests that they can occur in other species ^14,21^.

It is now well established that beyond their functions in the healthy brain, astrocytes play an active role in inflammatory responses associated with various pathological conditions, including neurodegenerative diseases ^22,23^. This activation encompasses a range of molecular, cellular, and functional alterations, typically characterized by increased expression of cytoplasmic proteins such as glial fibrillary acidic protein (GFAP) and vimentin, enlargement of the soma and cell processes, and disruption of their organized territorial domains ^24,25^. Consequently, the astrocytic response is dynamic and exists as a continuum that reflects the severity and progression of an underlying pathological condition ^26^. As aging, is the ground for many neurodegenerative processes, it is critical to understand how this condition modulates astrocytes in the brain.

Although evaluations of astrocyte changes during aging in rodents have provided a large amount of information, it is well known that primates, including humans have different astrocyte population. Evaluating astrocytes in different primates can thus provide a broader view of astrocytes evolution in aging. Studies in humans showed that aging is associated with a phenotypic shift, including upregulation of GFAP expression and morphological alterations, such as increased dendritic length, hypertrophy of the soma, and modifications of astrocytic territorial domains in different brain regions as the hippocampus and cortical regions ^25,27,28^. To the best of our knowledge, few studies reported white matter astrocytic changes in humans, except a study reporting an age-related reduction of astrocyte density in the white matter ^29^. In non-human primates such as macaques and chimpanzees, some studies suggested that age-related astrogliosis occurs mainly in white matter, with little change in cortical regions ^30,31^. However, other studies show that some aged chimpanzees exhibit cortical astrogliosis, with hypertrophic GFAP-positive astrocytes across all layers of the dorsolateral prefrontal cortex, sometimes accompanied by increased astrocyte density even without amyloid or tau pathology, the main lesions associated with Alzheimer pathology ^32^. It is thus critical to investigate astrocyte status in other primates to provide a larger perspective on astrocytes during aging.

The gray mouse lemur primate (*Microcebus murinus*) is a small primate (approximately 12 cm in length and 60–120 g in weight), endemic to Madagascar. It offers rodent-like practicality for primate research due to its rapid maturation (puberty at 6–8 months) and relatively short lifespan of approximately 12 years in captivity. Notably, a 10-year-old lemur is considered equivalent to a human centenarian, making it an ideal model for studying advanced aging ^33^. Research using the gray mouse lemur has yielded significant insights into cerebral aging and neurodegenerative diseases ^34,35^. For example, studies have shown a strong correlation between brain atrophy and cognitive deficits in older lemurs, paralleling similar phenomena in humans ^36^. Regarding the astrocyte characterization of this species, the interlaminar astrocytes have been well described by Colombo and coll.. as an intermediate stage between rodent and higher primate architecture ^37^. In a cohort of aged lemurs, the cerebral atrophy was associated to a reactive astrocytosis in the hippocampus ^38^.

Notably, no comprehensive studies have investigated astrocyte subtypes and their whole-brain distribution in the gray mouse lemur or their age-related changes. Here, we characterized GFAP- and vimentin-immunostained astrocytes across the entire brain of 17 lemurs, examining age-related changes. Astrocytes were predominantly localized in the white matter, with cortical astrocytes sparse in the parenchyma. Notably, pial and subpial ILAs bordering the pia and protoplasmic astrocytes along the white matter boundary formed prominent populations. Varicose projection astrocytes were observed near the white matter and in the hippocampus. The varicose aspect appeared as a shared phenotype across various radial glia-like cells, including tanycytes, border astrocytes of the basal hypothalamus, and ILAs. Aging primarily affected white matter astrocytes, increasing their density and size, and ILAs that exhibited denser palisades in older animals.

## 2. Materials and methods

### 2.1. Mouse lemurs

A total of seventeen mouse lemurs were studied with an age range of 1.0 to 11.5 years old (mean±SD: 8.8 ± 1.1 years) comprising four animals in middle-age (n=3 males, n=1 female, 3.1 ± 2.0 years old), and 13 aged animals (n=6 males, 7 females, 10.4 ± 1.5 years old; Supplementary table 1). The design and reporting of animal experiments were based on the ARRIVE reporting guidelines ^39^. The animals were reproduced in an approved breeding center (UMR 7179 CNRS/MNHN, France; European Institutions Agreement #962773) and provided to the animal facility of the Molecular Imaging Research Center, CEA, Fontenay-aux-Roses, where they were bred for the study. Mouse lemurs were kept in an enriched environment, temperatures between 24-26°C, relative humidity of 55%, and seasonal illumination (summer: 14h of light, 10h of darkness; winter: 10h of light, 14h of darkness). Food consisted of fresh apples and a handmade blend of bananas, cereals, eggs, and milk. Water supply for animals was freely accessible. None of the animals had ever taken part in invasive research or pharmaceutical trials. The 13 old animals and one middle-aged animal were sacrificed specifically for the study without presenting any identified pathology expected to lead to imminent spontaneous death. One middle-aged animal was euthanised following tail infection to shorten animal suffering. The brains of two middle-aged animals were sampled shortly after their spontaneous death from accident and sudden death. Magnetic resonance imaging was performed on all animals to rule out obvious cerebral pathology. MRI scans did not reveal any vascular pathology, severe atrophy process, or other cerebral impairment. Post-mortem autopsy was performed on each animal to evaluate hidden pathologies. Finally, histological staining for amyloid-β (Aβ) and Tau was performed on all the brains to rule out Alzheimer like pathologies (protocol detailed in ^35^). No peripheral chronic diseases were detected, and only one animal exhibited amyloidosis; this individual was excluded from the cohort. Due to the exploratory design of the study, the sample size was not defined in advance.

### 2.2. ETHICS

All experimental procedures were performed in compliance with the European Union directive on the protection of animals used for scientific purposes (2010/63/EU) and French regulations (Rural Code R214/87-131). They were approved by a local ethic committee (CETEA-CEA DSV IdF) as well as by the French Ministry of Education and Research (authorization A14_035 and A17_083). All efforts were made to minimize animal suffering and animal care was supervised by veterinarians and animal technicians skilled in mouse lemur healthcare and housing.

### 2.3. Perfusion and tissue preparation

The animals were killed with an intraperitoneal injection of pentobarbital sodium (0.1ml/100g; Exagon, Axience). Pentobarbital sodium is both an anaesthetic and a euthanasic agent. Here it is used as a euthanasic agent, but the death is preceded by an anaesthesia. Twenty minutes before any incision, a subcutaneous administration of buprenorphine (0.1mg/100g; Vétergésic®) was performed for analgesia. Animals were perfused intracardiacally with 0.1M PBS. The brain was post-fixed in 4% paraformaldehyde for 48h at 4°C., transferred in a 15% sucrose solution for 24h and in a 30% sucrose solution for 48h at 4°C for cryoprotection. Serial coronal sections of 40µm were performed with a microtome (SM2400, Leica Microsystem) and stored at -20°C in a storing solution (glycerol 30%, ethylene glycol 30%, distilled water 30%, phosphate buffer 10%). Free-floating sections were rinsed in a 0.1M PBS solution (10% Sigma-Aldrich® phosphate buffer, 0.9% Sigma-Aldrich® NaCl, distilled water) before use.

### 2.4. Immunohistochemistry

All tissues were incubated in hydrogen peroxide H2O2 30% (Sigma-Aldrich®) diluted 1/100 for 20 min to inhibit endogenous peroxidases. Blocking of non-specific antigenic sites was achieved over 30 min using a 0.2% Triton X-100/0.1M PBS (Sigma-Aldrich®) (PBST) solution containing 4.5% normal goat serum for GFAP protocol or 5% BSA for vimentin protocol. Sections were then incubated at +4°C GFAP (Dako Z0334, 1/10000) antibody diluted in a 3%NGS/PBST solution for 48h or vimentin (Dako MO725 1/1000) antibody diluted in a 3% BSA/PBST solution for 48h. After rinsing, an incubation with the appropriate biotinylated secondary antibody diluted to 1/1000 in PBST was performed for 1h at room temperature, followed by a 1h incubation at room temperature with a 1:250 dilution of an avidin-biotin complex solution (ABC Vectastain kit, Vector Laboratories®). Revelation was performed using the DAB Peroxidase Substrate Kit (DAB SK4100 kit, Vector 306 Laboratories®) with nickel sulfate. Sections were mounted on Superfrost Plus slides (Thermo-Scientific®). A cresyl violet counterstain was performed. All sections were then dehydrated in successive baths of ethanol at 50°, 70°, 96° and 100° and in xylene. Slides were mounted with the Eukitt® mounting medium (Sigma-Aldrich®).

### 2.5. Image acquisition and quantification

Stained sections were scanned using an Axio Scan.Z1 (Zeiss® - Z-stack images acquired at 20× (z-stacks with 16 planes, 1μm steps with extended depth of focus)). Each section was extracted individually in the .czi format using the Zen 2.0 (Zeiss®) software. Images were then imported in QuPath v0.4.3 software (the University of Edinburgh, UK (Bankhead et al., 2017)). Anatomical region of interest-hippocampus, corpus callosum and cortex - were manually segmented based on a mouse lemur atlas ^40^. A consistent positive threshold for DAB signal was then applied across all animals to quantify GFAP and vimentin staining in four coronal sections per animal. For vimentin analyses, one animal had to be excluded due to tissue damage. Astrocyte density in the white matter was determined by manual cell counting within three distinct regions of interest (ROIs) localized throughout the corpus callosum. In addition, astrocyte size in the white matter was estimated by dividing the GFAP-positive surface area by the corresponding manually counted astrocyte number, yielding an average GFAP-labeled area per astrocyte. Indexes of ILA extension were manually calculated in the parietal cortex using QuPath, by assessing the number of processes extending crossing virtual lines localized at 70 and 140 µm from cortical border (approximating crossing from cortical layer I to layer II and from layers II to III). Measures were performed on two different sections and encompassed a distance of 4mm after the interhemispheric fissure. Thus, the indexes of ILA extension represent the number of ILA processes per millimeter, calculated by dividing the total number of processes counted over the 4 mm distance by 4.

### 2.6. Statistical analysis

All statistical analyses were conducted using RStudio (RStudio Team, 2020) and R version 4.4.2. Age-related differences were assessed using pairwise permutation t-tests (pairwise.t.test in R) with 5000 permutations and Benjamini-Hochberg correction for comparisons across multiple regions. No prior ANOVA was performed, as analyses focused on independent, region-specific group differences. For single measures in a single region, the Exact Two-Sample Fisher-Pitman Permutation Test (Rcmdr package) was applied. Statistical significance was set at p < 0.05.

## Results

### 3.1. Astrocyte regional and morphological diversity in the mouse lemur brain

#### 3.1.1. Regional distribution and morphological features

In this study, we performed GFAP and vimentin immunostaining on a large population of gray mouse lemurs to characterize for the first-time astrocyte distribution at the whole brain level. Immunoreactive astrocytes were predominantly localized in white matter regions (Fig. 1a, j, k), including the corpus callosum (Fig. 1a, k), optic track (Fig. 1j) as well as in the hippocampus (Fig. 1f, h, l, o).

**Figure 1.**
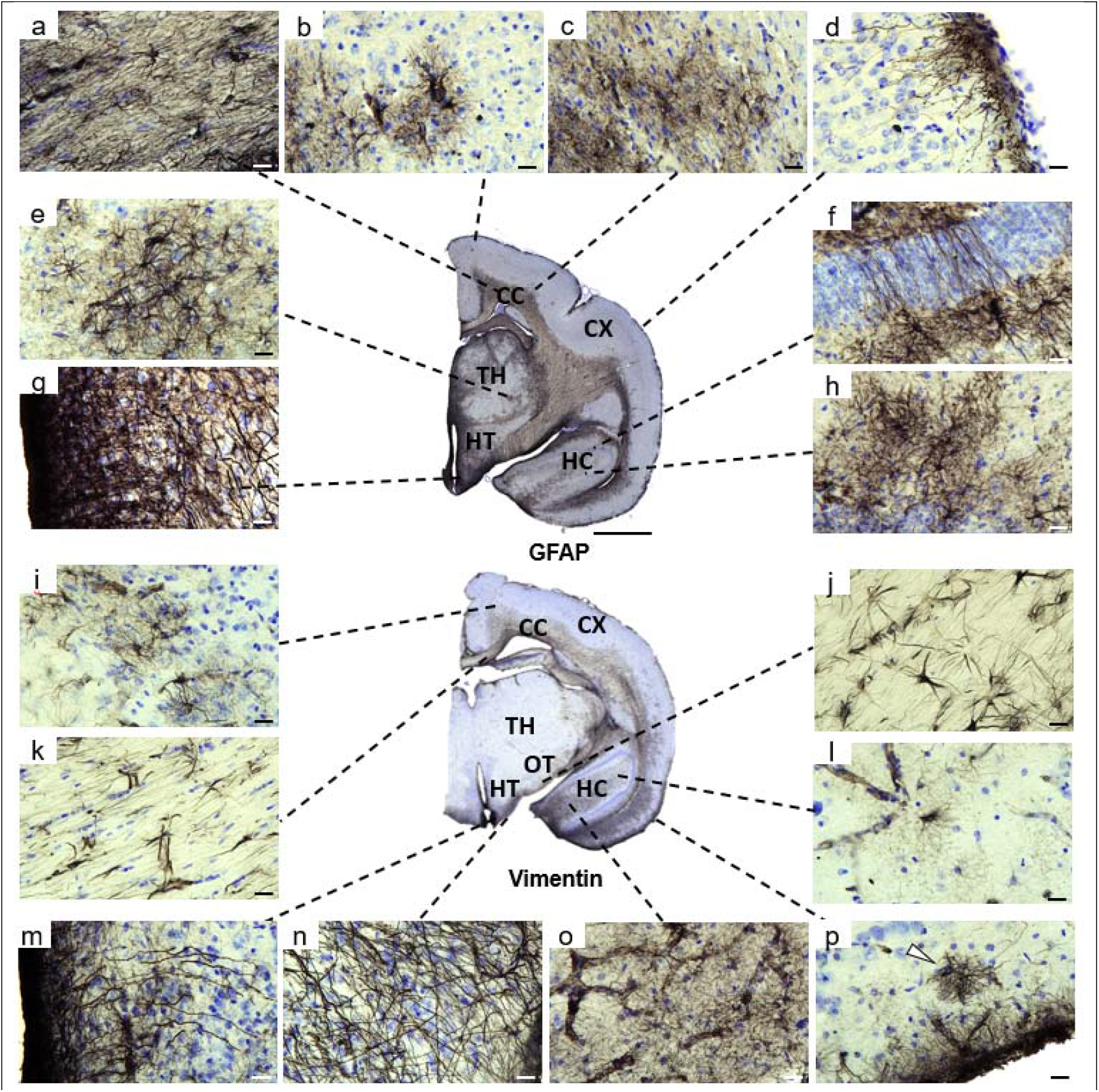
Distribution of GFAP and vimentin labeling in mouse lemurs. Most GFAP-expressing astrocytes (a–h) and vimentin-expressing astrocytes (i–p) were located in the white matter including corpus callosum (a, k) and optic track (j), whereas few cortical astrocytes were observed in the cortical parenchyma (b). The majority of cortical astrocytes were located in deep cortical layers near the corpus callosum (c, i) as well as along the pial surface, where subpial astrocytes (p, white arrow) and interlaminar astrocytes were observed (d, p). In the hippocampus, astrocytes form a syncytium-like network (h, l, o), with some polarized cells crossing the neuronal layer (f). GFAP+ astrocytes were present in the thalamus (e), whereas vimentin+ astrocytes were largely absent. In the hypothalamus, astrocytes with elongated processes extend from the pial surface to the parenchyma at the border of the basal hypothalamus and hypothalamic tanycytes display long processes surrounding the third ventricle (g, m). Scale bars: 20 µm (a–p) and 2 mm (tissue section). Abbreviations: corpus callosum (CC), hippocampus (HC), cortex (Cx), optic track (OP), thalamus (TH), hypothalamus (HT). Of note, the central section is representative and all the insets are not issued from this section. All images were obtained from old mouse lemurs.

In the white matter, fibrous astrocytes were regularly spaced from each other forming an interlocking network by extending elongated processes parallel to myelin fibers (Fig. 2a-c). These astrocytes varied in size and shape, ranging from large astrocytes with thick, elongated cell bodies and processes (Fig. 2a) to small star-shaped astrocytes with few processes (Fig. 2b). In addition, astrocytes were frequently observed in close association with blood vessels (Fig. 2c).

**Figure 2.**
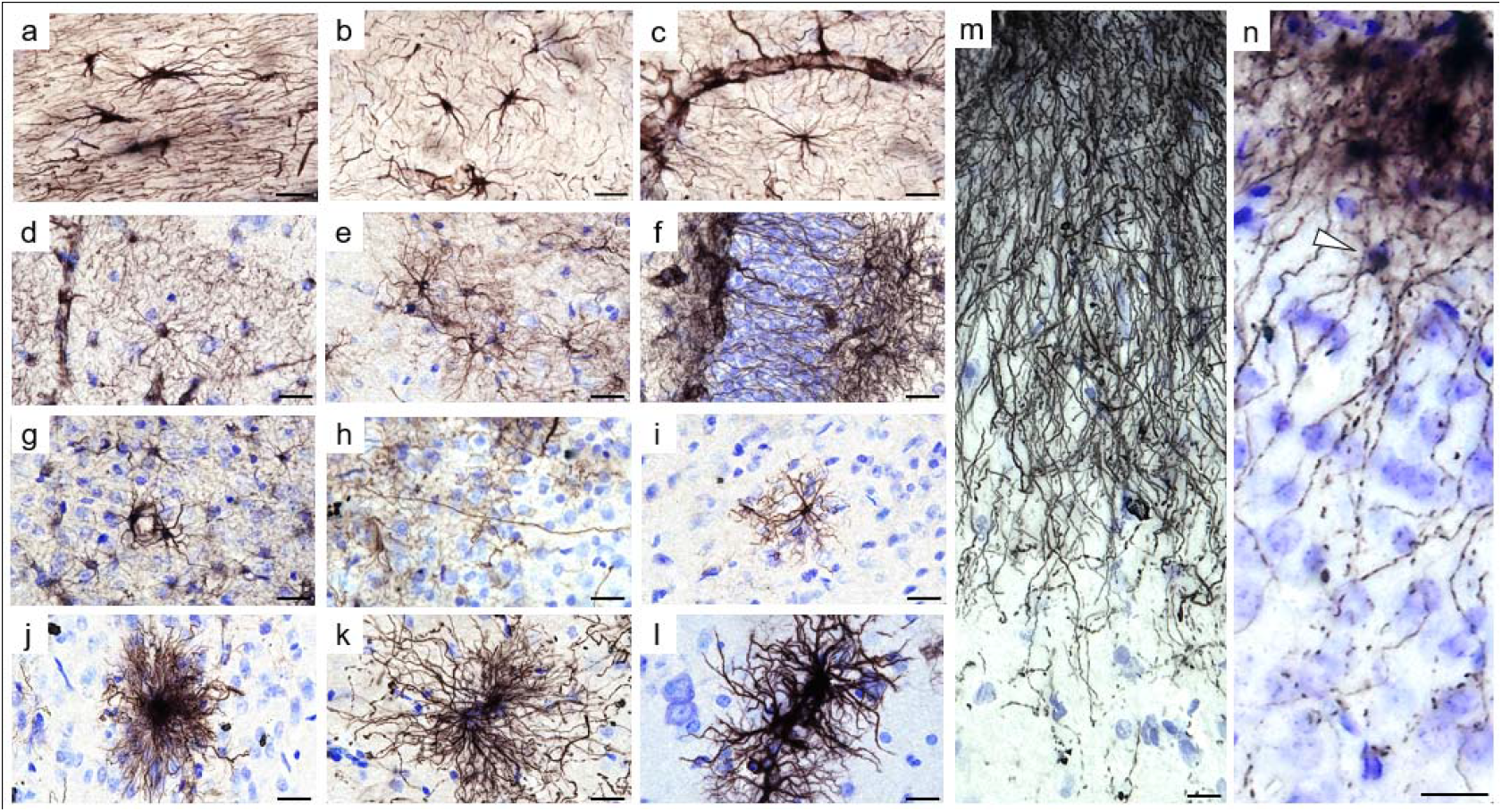
Astrocytes diversity in mouse lemurs. Fibrous astrocytes in the white matter displayed elongated processes aligned with myelin fibers (a) or a star-shaped morphology with few processes (b), often associated with blood vessels (c). Protoplasmic astrocytes in the hippocampus exhibited highly branched processes forming a dense network (d, e), with some processes crossing the neuronal layer (f). Parenchymal cortical astrocytes (g–i) were sparse and heterogeneous in size and morphology, including small astrocytes (i), contacting blood vessels (g), polarized astrocytes with long, thin unbranched process extending toward the upper layers (h), and giant, highly ramified astrocytes with bushy appearance or hypertrophied processes (j–l). At the pial surface, interlaminar astrocytes extended elongated processes reaching deeper cortical layers (m). A subpial interlaminar astrocytes with its soma (white arrow) located beneath the pia (n). Scale bar: 20 µm. Panels (d-f, l) are from middle-aged animals, whereas panels (a-c, g-k, m, n) are from old animals.

In the hippocampus, protoplasmic astrocytes were characterized by small round somata giving rise to numerous highly branched processes that formed a dense spongiform morphology. Each astrocyte occupied a distinct, non-overlapping territorial domain, contacting its neighbors only through the finest processes (Fig. 2d, e). Despite this spatial segregation, astrocytes were extensively interconnected, giving rise to a functional syncytium-like organization. We also observed polarized astrocytes with elongated processes that crossed the granule cell layer, thereby spanning multiple hippocampal laminae (Fig. 1f and Fig. 2f).

Another prominent astrocyte subtype in the mouse lemur was interlaminar astrocytes (ILAs), located at the pial surface of the upper cortical layers i.e. layer I (Fig. 1d, p, Fig. 2m-n). These astrocytes were in contact with the pia mater and extended long, parallel processes into deeper cortical layers, forming a characteristic palisade-like pattern (Fig. 2m). Of note, ILAs with long processes extending beyond layer I (Fig. 2m) are referred to as “typical” ILAs ^41^. We also detected ILAs with short processes that remain confined to layer I (Fig. 1p). They are referred to as “rudimentary” ILAs ^41^. Among ILAs, a subset of subpial ILAs was observed, with the soma located within layer I and processes oriented both toward the pia mater and toward deeper cortical layers (Fig. 2n, white arrow). Subpial ILAs remain rare, and most interlaminar processes emerge from pial ILAs, with only a few processes arising from subpial ILAs. In addition, some astrocytes were observed in layer I along the pial surface, corresponding to subpial protoplasmic astrocytes (Fig. 1p-white arrow).

Other cortical regions, in contrast to the white matter and hippocampus, exhibited sparse labeling (Fig. 1b–d, i, p), with only a few isolated astrocytes observed in the parenchyma (Fig. 1b). Most were located in deep cortical layers near the white matter, often in close proximity to each other, contacting blood vessels (Fig. 2g). Some of the rare astrocytes observed in deep cortical layers, exhibited a polarized morphology, characterized by a long unbranched process extending towards upper layers (Fig. 2h) resembling polarized astrocytes described in the human cortex ^19^ as well as in ferrets ^14^. The other cortical astrocytes were heterogeneously distributed and morphologically diverse, ranging from small protoplasmic astrocytes (Fig. 2i) to large cells with highly ramified processes (Fig. 2j) or hypertrophic features, including thickened processes and enlarged domains (Fig. 2k, l).

In the thalamus and basal hypothalamus, GFAP-immunoreactive astrocytes were abundant, whereas vimentin staining was low or largely restricted to endothelial cells (Fig. 1e). A high density of radial glia-like astrocytes (both GFAP and vimentin-positive) was observed along the ventral border of the basal hypothalamus (Fig. 1n). In addition, specialized radial glia known as tanycytes (considered here as a subcategory of astrocytes) were present around the third ventricle (Fig. 1g).

#### 3.1.2. Widespread varicosities across astrocyte subtypes

Notably, many astrocytic processes throughout the brain, including those in the cortex, hippocampus, and hypothalamus, exhibited bead-like structures known as varicosities. Varicose projection astrocytes (VPAs) were observed in the parenchyma, including the hippocampal granule cell layer, at the hippocampal–corpus callosum border, predominantly in cortical layers IV–V near the corpus callosum (Fig. 3b–e), and in the hypothalamus parallel to the wall of the third ventricle (Fig. 4b). Varicosities were also present in astrocytes with a radial shape and localized close to brain borders, such as ILAs at the cortical pia (Fig. 3a), and in tanycytes bordering the third ventricle (Fig. 4c). In addition, astrocytes lining the basal border of the hypothalamus, displayed a varicose phenotype in their elongated and polarized processes (Fig. 4d). In all these astrocytes, two types of varicosities were observed: continuous varicosities connected along the processes (Fig. 3a’), and interrupted varicosities giving a fragmented appearance (Fig. 3a’’). Collectively, these observations indicate that varicosities are a widespread morphological feature across multiple astrocytic subtypes in the gray mouse lemur, spanning cortical, hippocampal, and hypothalamic regions.

**Figure 3.**
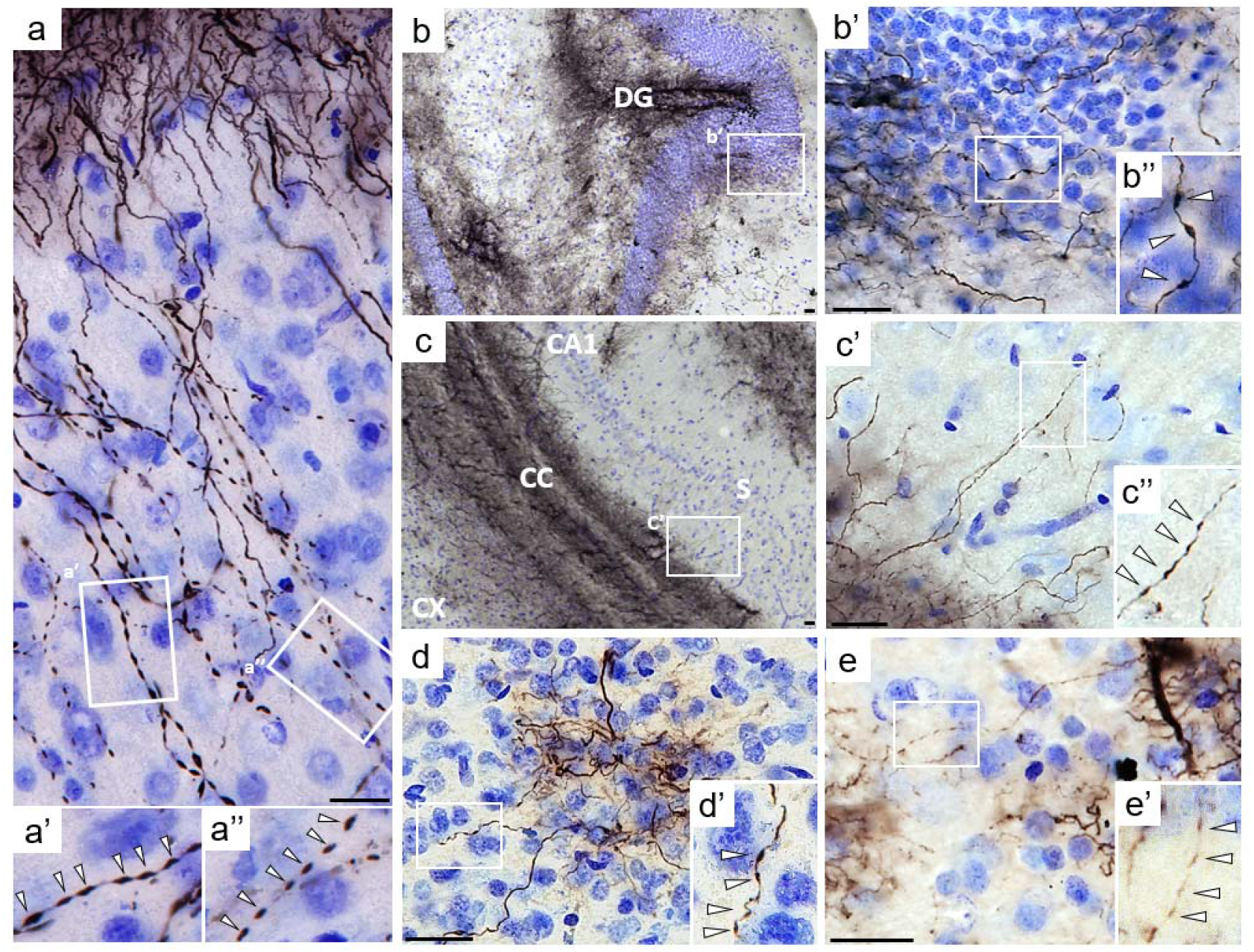
Varicosities across different astrocyte subtypes. Varicosities in interlaminar astrocytes in the superficial cortex (a–a’’), with higher magnifications (a’, a’’) showing continuous and fragmented varicosities. Varicose projection astrocytes (VPAs) in the dentate gyrus (DG) granular layer (b–b’’), in the hippocampal subiculum (S) at the border with the corpus callosum(CC) (c-c’’), in cortical layers IV (d–d’), and in cortical layer V at the corpus callosum border (e–e’); zoomed panels highlight varicosities (white arrows in a’, a’’, b’’, c’’, d’, e’). Scale bars: 20 µm. Panel (a) is from a middle-aged animal, whereas panels (b–e) are from old animals. Abbreviations: CX, cortex; CC, corpus callosum; DG, dentate gyrus; CA1, cornu Ammonis 1; S, subiculum.

**Figure 4.**
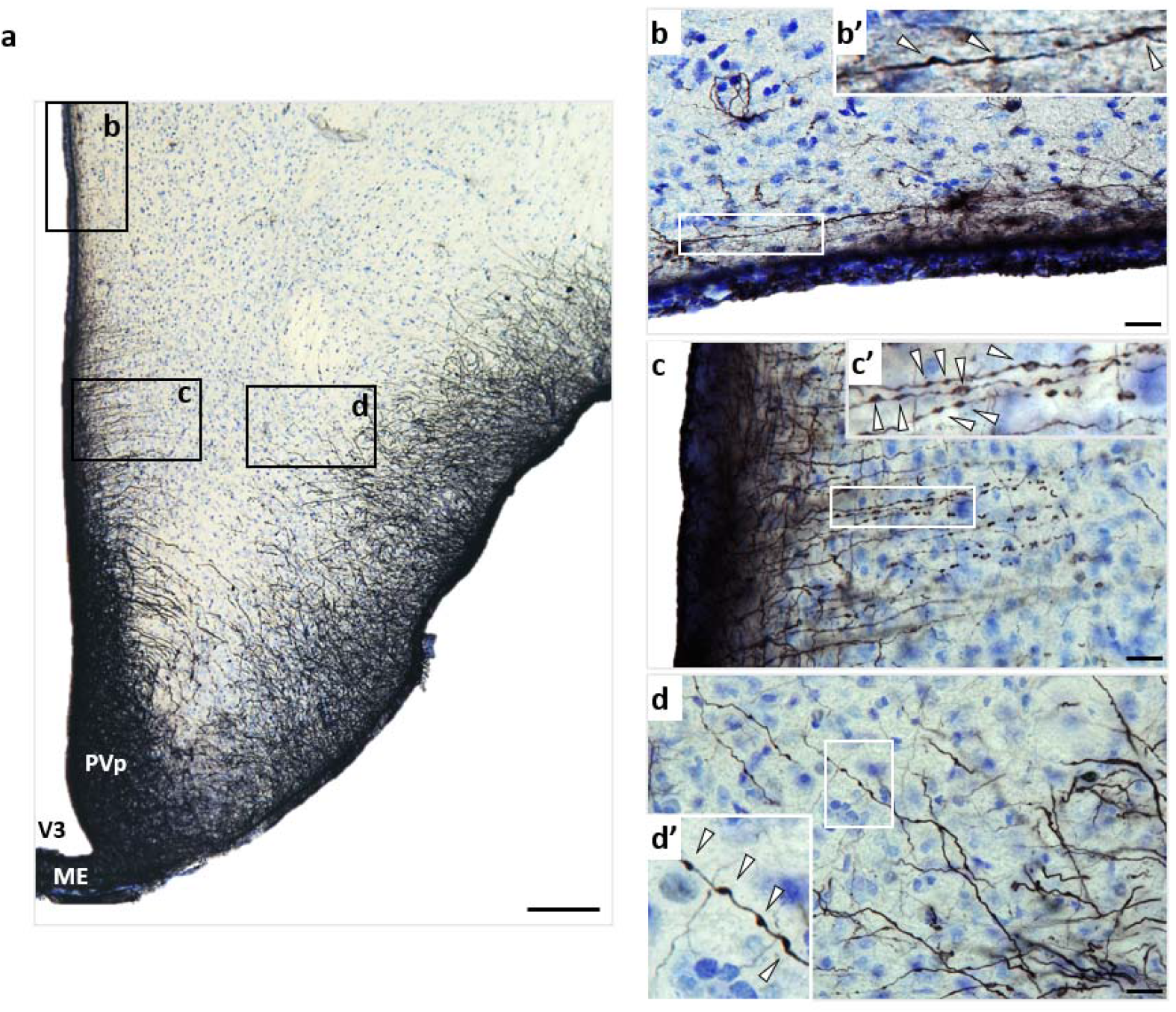
Varicose phenotypes in hypothalamic glial cells (vimentin immunostaining) Basal hypothalamus of an old mouse lemur (a, scale bar: 100 µm). A varicose projection astrocyte (VPA) located at the border of the third ventricle (b, scale bar: 20 µm). Tanycytes along the walls (PVp) and floor (ME) of the third ventricle (V3), display bead-like structures resembling varicosities (c, scale bar: 20 µm). Astrocytes with elongated processes lining the basal border of the hypothalamus, exhibiting a varicose phenotype (d, scale bar: 20 µm). Higher-magnification views of the boxed areas are shown, with white arrows indicating individual varicosities (b’, c’, d’). Abbreviations: ME = median eminence, PVp = periventricular hypothalamic nucleus, posterior part, V3 = third ventricle.

### 3.2. Age-dependent regional astrocyte alterations in the mouse lemur brain

#### 3.2.1. Age-related increases of astrocyte density and hypertrophy in the white matter

Visual observation of age-related immunoreactivity revealed considerable inter-individual variability in white matter areas. Middle-aged animals showed generally low staining (Fig. 5a-b, column 1), whereas older animals displayed a wide range of profiles, from moderate (Fig. 5a-b, column 2) to very high levels of GFAP or vimentin expression (Fig. 5a-b, column 3). Notably, some older individuals exhibited low immunoreactivity, comparable to that of middle-aged animals, indicating that aging was associated with highly heterogeneous astrocytic responses, ranging from profiles of pronounced astrogliosis to profiles with minimal changes. Overall, GFAP expression (Fig. 5a) remained stronger than vimentin (Fig. 5b) across all ages. We then quantified astrogliosis by evaluating astrocyte global immunoreactivity as well as individual astrocyte densities and sizes. Quantitative analysis highlighted a significant increase in astrocyte immunoreactivity within the white matter of old animals compared to middle-aged ones (GFAP: *42% positive area in middle aged versus 66% in old lemurs*, p = 0.03; vimentin: *7.9% for the middle aged versus 25% in old lemurs* p =0.049, Fig. 5c-d). GFAP and vimentin-IR in the white matter were highly correlated (data not shown).

**Figure 5.**
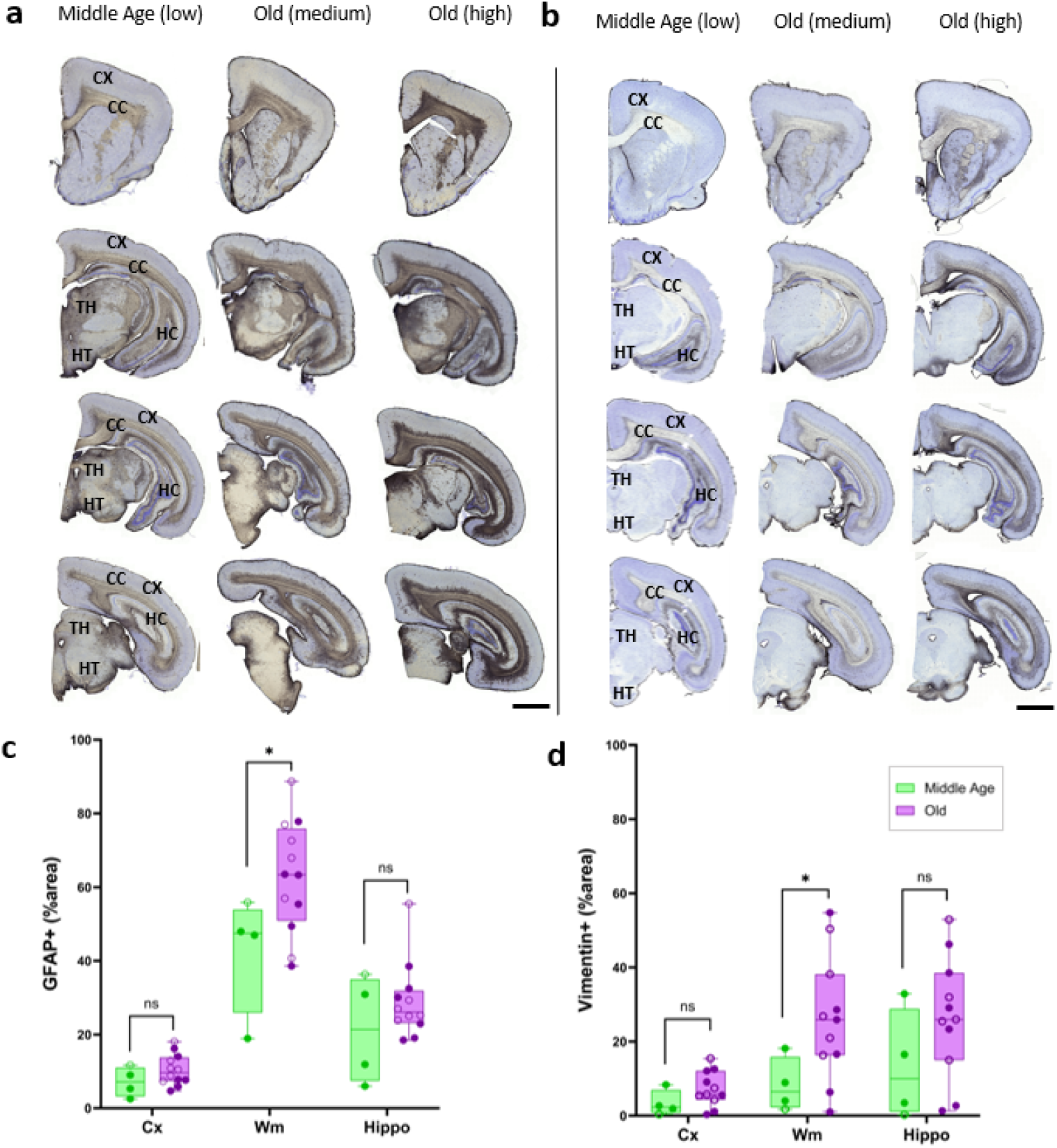
Age-related astrocytes immunoreactive profiles in mouse lemur brain. GFAP (a) and vimentin (b) astrocyte immunoreactivity (from frontal to posterior brain sections) is low in middle-aged brains (column 1), moderate and high in old lemurs (columns 2 and 3). Abbreviations: corpus callosum (CC), hippocampus (HC), cortex (Cx), thalamus (TH), hypothalamus (HT). Significant age-related increase of GFAP (c) and vimentin (d) immunoreactivity (% area) in the cortex (Cx), white matter (Wm), and hippocampus (Hippo) of old (n = 12) compared with middle-aged (n = 4) lemurs. Filled symbols denote males, and open symbols denote females. Statistical analysis was performed using R (version 4.3-3) with pairwise.perm.t.test (1000 permutations, BH correction for multiple comparisons); *p < 0.05.

To further investigate age-related white matter changes, we measured astrocyte density in the corpus callosum. In aged animals, astrocyte density was higher for both GFAP+ (p = 0.013, Fig. 6b) and vimentin+ (p = 0.039, Fig. 6d) astrocytes compared to middle-aged animals (Fig. 6a, c). Visual observation outlined that astrocytes were hypertrophic exhibiting a thicker cell body and more robust processes in aged animals compared to middle age ones (Fig. 6b, d versus Fig. 6a, c). To quantify these morphological changes, we estimated astrocyte size by dividing the total positive immunoreactive surface area by the number of astrocytes within the white matter region of interest. This analysis confirmed a significant increase in GFAP+ astrocyte size in old animals (p = 0.04, Fig. 6f) and a trend toward increased vimentin+ astrocyte size (p = 0.08, Fig. 6h). Furthermore, astrocyte density and size were strongly correlated for both GFAP+ (r=0.62, p=0.008, Fig. 6i) and Vim+ (r=0.82, p=0.0001, Fig. 6j) astrocytes.

**Figure 6.**
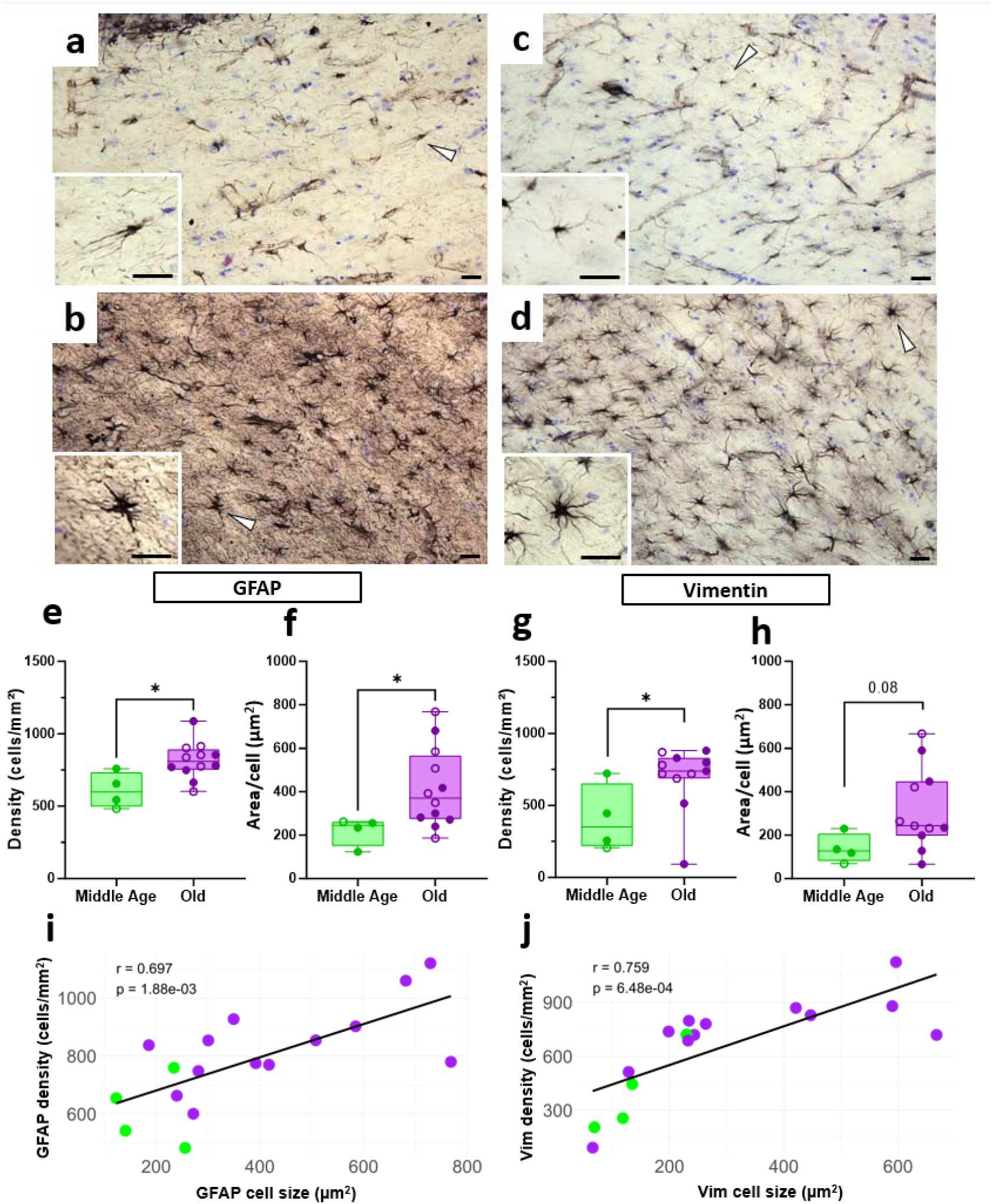
Age-related astrocyte reactivity in the white matter of mouse lemur. GFAP-immunoreactive astrocytes in the WM of a middle-aged (a) and an old lemur (b). Vimentin-immunoreactive astrocytes in the WM of a middle-aged (c) and an old lemur (d). White arrows indicate the astrocyte shown at higher magnification. Scale bar: 25µm. Increased GFAP (e-f) and vimentin (g-h)) astrocyte density (cells/mm^2^, e, g) in the WM of middle-aged and old lemurs and astrocyte domain (surface area labeled /number of astrocytes, f, h) in the WM of old animals compared to middle-age lemurs. Filled symbols denote males, and open symbols denote females. Statistical analysis is performed using Rcmdr – R package version 4.3-3 with Exact Two-Sample Fisher-Pitman Permutation Test; *p < 0.05. Correlation between density and estimated cell size for GFAP (i) and vimentin immunoreactive astrocytes (j). Pearson Correlation test was used. Solid lines represent linear regression fits.

#### 3.2.2. Cortical astrocytes: Age-dependent changes in interlaminar astrocytes

A first analysis of astrocyte age-related change of immunoreactivity excluding astrocytes from border cortical areas did not detect age-related changes of astrocytes in the cortex and in the hippocampus (Fig. 5c-d). The cortical region used for this analysis excluded pial astrocytes to avoid potential bias from border effects. To complete this analysis, we investigated the interlaminar astrocyte (ILA) palisades separately. This structure appeared denser in localized patches along the pia. As shown in Fig. 7a-b, the ILA palisade was notably denser in aged animals. By quantifying the number of ILAs processes extending into cortical layers II and III in the parietal cortex, we observed a significant age-related increase in the number of GFAP+ ILAs processes reaching layer II (p = 0.03, Fig. 7c) and a strong tendency for layer III (p = 0.08, Fig. 7c). No significant age-related increase was observed for Vim+ ILAs processes (p > 0.05, data not shown).

**Figure 7.**
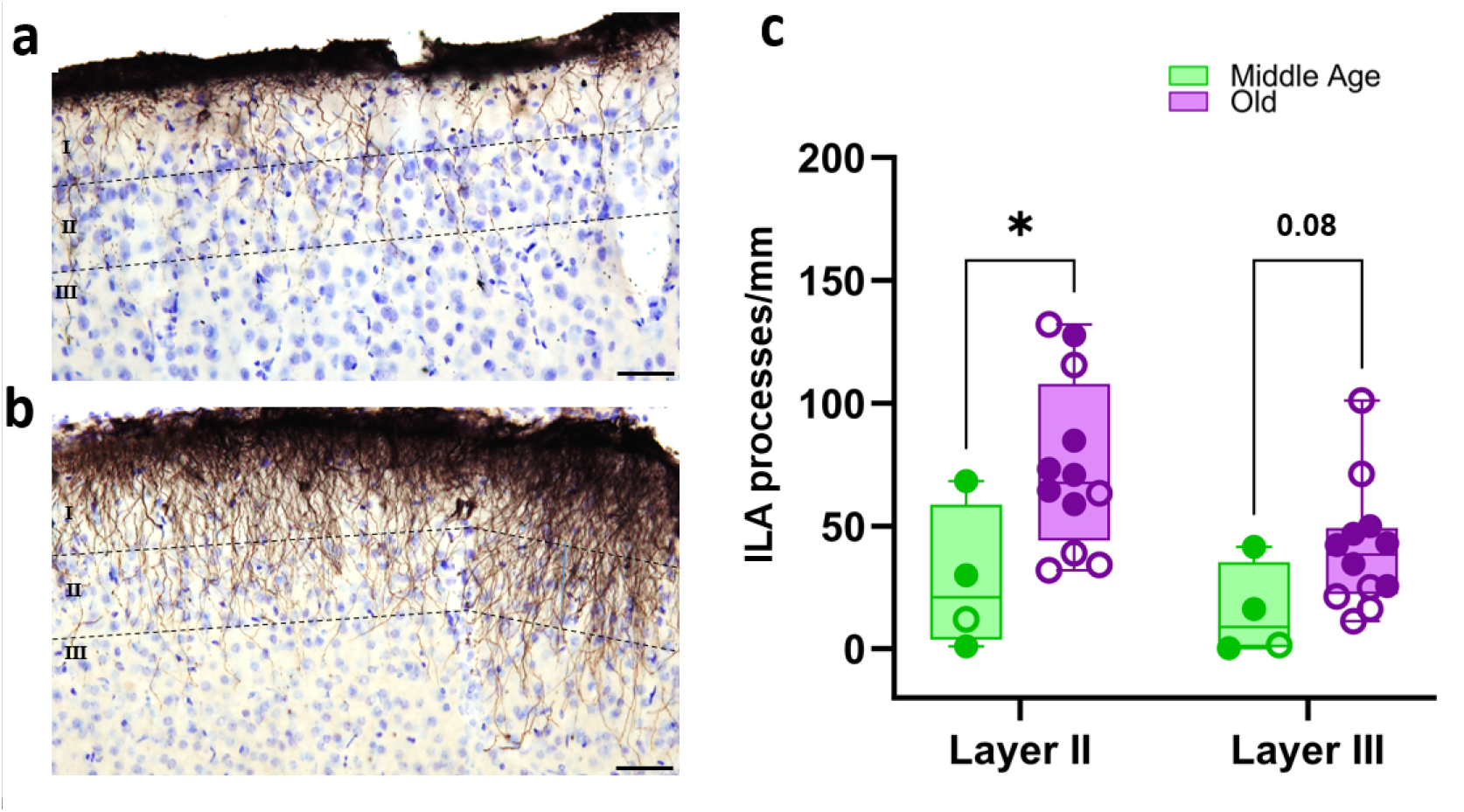
Age-related increase in interlaminar astrocytes (ILAs) processes. Representative images of the ILA palisade in the parietal cortex of a middle-aged (a) and an old (b) animal, stained with GFAP. Dashed lines approximately delineate cortical layers I-III. Scale bar: 50 µm. Quantification of GFAP+ ILAs processes crossing cortical layer II or III in the parietal cortex (c). Filled symbols denote males, and open symbols denote females. Statistical comparisons between groups were performed using the Rcmdr package (R version 4.3-3) with Exact Two-Sample Fisher-Pitman Permutation Test; *p < 0.05, ns, not significant (p > 0.05).

## 4. Discussion

### 4.1. Astrocyte subtype diversity in the mouse lemur brain

In this study, we provide the first comprehensive description of astrocyte distribution and subtypes in the gray mouse lemur, revealing a remarkable diversity of morphologies across brain regions. At first sight, astrocytes were mostly visible in white matter regions such as the corpus callosum and optic track. These astrocytes were fibrous astrocytes. Such astrocytes are arranged in networks, where they support oligodendrocytes by facilitating myelination during development and adulthood (for review see Lundgaard and coll. ^16^). Another striking feature detected in mouse lemurs is the presence of cortical ILAs, which are located at the pia level and represent a distinctive characteristic of primates or some carnivores ^14,42^. While the precise origin and function of ILAs remain subjects of debate, it is hypothesized that their columnar organization plays a key role in information trafficking between cortical layers, as well as in communication with capillaries, meninges, cerebrospinal fluid, and potential involvement in the glymphatic system ^13,41,43^. In addition to pial ILAs, we observed subpial ILAs, characterized by a soma located in the middle of layer I and long polarized processes extending both toward the pia mater and deeper cortical layers, a feature only described in primates ^41^. Two main categories of pial or subpial ILAs have been reported in the literature, referred to as “typical” and “rudimentary”. Typical ILAs have long processes that exit layer I, whereas rudimentary ILAs remain confined within layer I ^41^. Pial ILAs, are present in their typical or rudimentary forms across mammals. For instance, primates display typical ILAs while rodents exhibit only rudimentary ILAs. In carnivore order, mainly rudimentary pial and subpial ILAs with processes restricted to layer I are observed ^14^, although some typical ILAs have been reported in the cortical ventral regions only ^41^. In mouse lemurs, we identified both rudimentary and typical ILAs (ventral and dorsal), suggesting a cortical astroglial architecture that is intermediate between non-primate mammals and higher primates ^37^. These observations support the idea that rudimentary ILA may develop into typical ILA across mammalian evolution ^41^.

In other cortical areas, distant from layer I regions, few astrocytes were observed. The most abundant reactive astrocytes were in deep cortical layers near the white matter. Some of these cells extended long, thin unbranched processes that crossed multiple cortical layers. Such long-projections astrocytes were first described in humans as polarized astrocytes with varicosities along their processes, also referred to as varicose projection astrocytes (VPAs) ^19^. Here, we report for the first time the presence of polarized astrocytes in the mouse lemur. To our knowledge, we are the first to show these polarized astrocytes without varicosities. We also detected sparse protoplasmic astrocytes in the middle cortical layers and a few subpial protoplasmic astrocytes in the superficial cortical layer I.

Interestingly, in the cortex, some astrocytes did display varicosities along their processes consistent with the VPA morphology. VPAs were also observed in the hippocampus, either spanning the granular layer or located near the corpus callosum. In addition to their presence in projection astrocytes, varicosities were also detected in other subtypes of radial glia-like cells, including ILAs, tanycytes, and border astrocytes lining the basal hypothalamus. Notably, a study on human brain tissue reported an increased prevalence of VPAs in both aging and disease contexts ^21^, suggesting that this varicose phenotype represents a non-physiological state of astrocytes. VPAs were initially described as a unique feature of higher primates and humans ^20^, but more recent work demonstrated that they can be detected in many other species like in ferret ^14^ or tiger ^21^. Usually, varicosities are described as interconnected GFAP or vimentin-positive beads connected by thin processes. Here, we reported for the first time, non-connected beads, suggesting that varicosities may become interrupted. These observations raise the possibility that the varicose phenotype evolves through distinct stages, ultimately leading to fragmented processes.

### 4.2. Astrocyte reactivity and aging: similarities and differences across species

Aging is a well-known condition that modulates astrocytes. Here, we demonstrated that age-related astrocytic reactivity in the gray mouse lemur is highly region-specific, with white matter showing the most pronounced age-related astrogliosis, whereas the deep cortical region and hippocampus exhibit only modest changes. This aligns with studies in macaques that show predominantly age-related astrocytic increase in the white matter ^30,44^. Given the prominent changes in astrocytic reactivity in the white matter with aging in mouse lemurs, further studies will have to investigate interaction between astrocytes and white matter integrity in lemurs.

Interestingly, in mouse lemurs, we outlined that interlaminar astrocytes (ILAs) were the densest and most structurally prominent astrocytic cortical subtype. They formed palisade-like arrangements that extend processes from superficial cortical layer I across deeper cortical layers and became more prominent in aged-animals. This indicates that ILAs represent a key site of age-related astrocytic change in the lemur cortex. This observation parallels findings in chimpanzees ^32^, where cortical layer I astrocytes (corresponding to the location of ILAs) exhibit the most significant astrogliosis. In rhesus monkeys, the age-related thickening of the glial limiting membrane and the increased astrocytic filament density in layer I ^45^ also echo the rise in ILAs processes density observed during aging in the present study.

The modest astrocytic changes reported in other cortical areas in lemurs is consistent with data in macaques and chimpanzee, in which most cortical and hippocampal astrocytes remain largely stable with age unless challenged by pathology, such as amyloid or tau deposition ^30,32^.

Finally, we also observed considerable age-related inter-individual variability, with some aged animals exhibiting minimal astrocytic reactivity while others displayed pronounced astrogliosis, a pattern reminiscent of that described in aged chimpanzees, where astrocytic changes vary widely across individuals and brain regions ^32^. To the best of our knowledge this heterogeneity as not been investigated in humans. This point emphasizes the need to further consider aging trajectories at the level of astrocytes.

### 4.3. Conclusion

Overall, this study characterized seven different astrocyte subtypes: from fibrous to primate-specific astrocyte as interlaminar astrocytes in different brain regions of mouse lemurs. It underscores the heterogeneity of astrocyte aging across brain regions, emphasizing the particular vulnerability of the white matter and ILAS in the aging primate brain. The presence of varicosities across multiple astrocytic subtypes—including ILAs, VPAs, and hypothalamic astrocytes—further highlights a shared phenotype feature that may reflect adaptive or stress-related responses. Finally, the diversity of astrocytic subtypes and their phenotypic similarities to human astrocytes highlight the gray mouse lemur as a valuable translational model to investigate glial changes in aging and neurodegenerative diseases.

## 5. Acknowledgements

We thank the Agence Nationale de la Recherche (PrimAlz, ANR-22-CE14-0056), France-Alzheimer Association for funding this study. The MIRCen facility was funded by a grant from NeurATRIS: A Translational Research Infrastructure for Biotherapies in Neurosciences (“Investissements d’Avenir”, ANR-11-INBS-0011). LG was financed by the French Ministère de l’Enseignement Supérieure, de la Recherche et de l’Innovation.

## 8. Data and code availability statements

The data that support the findings of this study are available from the corresponding author upon reasonable request.

## 10. Author contributions

L.G, L.D, M.D contributed to the study conception and design. F.P and S.L performed the microcebe euthanasia and collected brain samples. L.G, F.P, S.L performed immunohistological experiments. L.G conducted quantification. L.G and L.D analyzed the data. L.G, L.D, J.L.P and M.D wrote the manuscript. M.D edited the manuscript. All authors commented on previous versions of the manuscript. All authors read and approved the final manuscript.

## 11. Supplementary data

**Supplementary table 1**

Characteristics of the gray mouse lemurs included in the study. Age, sex, and relevant medical information are reported; if no particular observation was noted, “NSR” (Nothing Special to Report) is indicated. FD: Found Dead; SE: Euthanized for study purposes; AE: Euthanized to prevent suffering due to a diagnosed pathology. Histological features for each animal are also provided, including the presence or absence of Alzheimer’s disease-related neuropathology, as well as white matter (WM) and gray matter (GM) astrocytic parameters measured in the corresponding regions. The animal with AD-related pathology was excluded from all statistical analyses to avoid bias.

**Supplementary table 2**

Key resource table

