## Supplementary data for "Astrocyte diversity and aging in the mouse lemur primate brain"

### Supplementary table 1

Characteristics of the gray mouse lemurs included in the study. Age, sex, and relevant medical information are reported; if no particular observation was noted, "NSR" (Nothing Special to Report) is indicated. FD: Found Dead; SE: Euthanized for study purposes; AE: Euthanized to prevent suffering due to a diagnosed pathology. Histological features for each animal are also provided, including the presence or absence of Alzheimer's disease-related neuropathology, as well as white matter (WM) and gray matter (GM) astrocytic parameters measured in the corresponding regions. The animal with AD-related pathology was excluded from all statistical analyses to avoid bias.

| | | | | | | WM Astrocytes | | | | | | GM Astrocytes | | | | | |
|---|---|---|---|---|---|---|---|---|---|---|---|---|---|---|---|---|---|
| | | | | | | GFAP | | | Vimentin | | | GFAP | | | Vimentin | | |
| | | | | | | corpus callosum | | | | | | cortex | | hippocampus | cortex | | hippocampus |
| Age | Sexe | Medical record | Death context | Aβ | Tau | surface area (% area) | density (cell/mm²) | size (µm²) | surface area (% area) | density (cell/mm²) | size (µm²) | surface area (% area) | linear density (ILA/mm) | surface area (% area) | surface area (% area) | linear density (ILA/mm) | surface area (% area) |
| 2.3 | M | Sudden death | FD | - | - | 55.9 | 542.9 | 141.0 | 4.0 | 256.7 | 118.5 | 9.0 | 30.0 | 36.3 | 1.8 | 3.0 | 3.4 |
| 4.1 | M | NSR | SE | - | - | 47.9 | 759.9 | 234.4 | 18.1 | 721.8 | 229.8 | 7.6 | 68.3 | 30.9 | 8.3 | 67.3 | 32.9 |
| 4.8 | M | Tail infection | AE | - | - | 18.9 | 654.8 | 123.1 | 8.9 | 445.4 | 134.9 | 7.8 | 1.0 | 11.9 | 2.6 | 0.0 | 16.5 |
| 5.3 | F | Head Shock | SE | - | - | 47.0 | 483.5 | 256.0 | 1.7 | 205.7 | 68.8 | 8.9 | 12.0 | 6.0 | 0.4 | 2.5 | 0.2 |
| 9.7 | M | Cataract | SE | - | - | 49.4 | 748.6 | 281.5 | 25.9 | 798.5 | 234.1 | 10.9 | 73.2 | 23.9 | 9.0 | 37.5 | 38.5 |
| 9.8 | F | NSR | SE | - | - | 40.7 | 838.0 | 186.1 | 16.3 | 719.8 | 243.4 | 7.4 | 34.3 | 22.9 | 4.3 | 21.3 | 15.0 |
| 9.8 | F | NSR | SE | - | - | 72.6 | 927.7 | 349.6 | 38.1 | 869.9 | 421.2 | 14.0 | 115.5 | 27.0 | 7.4 | 88.8 | 25.4 |
| 9.8 | M | NSR | SE | - | - | 38.6 | 663.5 | 240.1 | 16.5 | 738.7 | 198.6 | 12.9 | 127.8 | 19.1 | 12.0 | 5.5 | 46.2 |
| 10.2 | M | Cataract | SE | - | - | 63.3 | 770.2 | 417.8 | 28.6 | 829.4 | 447.1 | 4.7 | 84.8 | 30.1 | 5.5 | 76.3 | 29.1 |
| 10.2 | M | Eye ulcer | SE | - | - | 57.0 | 854.1 | 300.7 | 1.0 | 92.4 | 65.7 | 18.1 | 70.8 | 18.5 | 0.3 | 3.5 | 1.3 |
| 10.2 | F | NSR | SE | - | - | 55.4 | 601.2 | 271.9 | 20.9 | 687.9 | 232.5 | 2.6 | 39.0 | 29.3 | 5.4 | 25.8 | 31.9 |
| 10.2 | F | NSR | SE | - | - | 77.0 | 854.5 | 507.9 | 50.4 | 719.4 | 667.4 | 11.7 | 132.0 | 32.5 | 15.4 | 94.8 | 52.9 |
| 10.5 | M | NSR | SE | - | - | 77.8 | 1060.2 | 681.6 | 54.8 | 879.7 | 590.0 | 10.5 | 59.0 | 25.0 | 12.6 | 42.3 | 23.3 |
| 10.5 | F | NSR | SE | - | - | 67.9 | 775.6 | 392.1 | 26.8 | 781.1 | 263.6 | 5.3 | 63.3 | 25.2 | 5.7 | 25.3 | 25.9 |
| 10.9 | M | NSR | SE | - | - | 63.5 | 780.3 | 768.5 | 6.3 | 513.6 | 128.6 | 16.2 | 64.5 | 38.5 | 1.1 | 6.5 | 2.7 |
| 11.3 | F | NSR | SE | - | - | 88.7 | 902.9 | 584.7 | NA | 675.0 | NA | 11.8 | 32.0 | 55.5 | NA | NA | NA |
| 11.4 | F | Cataract | SE | + | - | 89.9 | 1120.3 | 729.0 | 64.0 | 1124.7 | 596.5 | 5.9 | 0.3 | 26.1 | 2.7 | 0.0 | 7.0 |

### Supplementary table 2

Key resource table

| Reagent or Resource | Source | Identifier |
|---|---|---|
| **Antibodies** | | |
| Rabbit polyclonal anti-GFAP<br>Dilution: 1/10000 | Dako | Cat#Z0334 |
| Mouse monoclonal anti-Vimentin<br>Dilution: 1/1000 | Dako | Cat#M0725 |
| HRP Goat Anti Rabbit IgG | ThermoFisher | Cat#31460 |
| HRP Goat Anti Mouse IgG | ThermoFisher | Cat#31430 |
| **Chemicals and commercial assay or kit** | | |
| ABC Vectastain® ABC-HRP kit | Vector Laboratories® | Cat#PK6100 |
| Bovine serum albumin (BSA) | Sigma-Aldrich® | Cat#A7906 |
| Normal goat serum (NGS) | Sigma | Cat#G6767 |

| | | |
|---|---|---|
| Cresyl violet | Merck | Cat#10510-54-0 |
| DAB Substrate kit, Peroxidase (HRP), with Nickel | Vector Laboratories® | Cat#SK4100 |
| Dulbecco's phosphate saline (DPBS) 1X | Gibco™, ThermoFisher | Cat#14190094 |
| Ethanol absolute | VWR | Cat#83813360 |
| Ethylene glycol | Carlo Erba | Cat#346502 |
| Eukitt® mounting medium | Sigma-Aldrich® | Cat#03989 |
| Glycerol | Fisher | Cat#12144481 |
| Hydrogen peroxide 30% | Sigma-Aldrich® | Cat#H1009 |
| Paraformaldehyde, PFA | Sigma | Cat#P7148 |
| Buprenorphine | Vétergésic® | |
| Pentobarbital | Exagon®, Axience | |
| Phosphate buffer solution, 1 M , pH 7.4 | Sigma-Aldrich® | Cat#P3619 |
| Phosphate Buffered Saline (PBS), pH 7.4 | Sigma-Aldrich® | Cat#806552 |
| Sodium chloride (NaCl) | Sigma-Aldrich® | Cat#S9888 |
| Sucrose | Sigma-Aldrich® | Cat#S0389 |
| Tris-HCl | Merck | Cat#252859-500G |
| Triton X-100 | Sigma-Aldrich® | Cat#X100 |
| Xylene | VWR Chemicals | Cat#28973363 |
| **Equipements** | | |
| Camera | JENOPTIK GRYPHAX® | |
| Axio Scan.Z1 | Zeiss® | |
| Microscope | Leica DMI6000 | |
| Microtome | Leica Vt1200 blade | |
| Perfusion pump | Fisher Scientific | Cat#1170-5369 |
| SM2400 microtome | Leica Microsystems | |
| **Experimental models: Organisms / Strains** | | |
| Microcebus Murinus (WT) | Brunoy - MNHN | |
| **Material** | | |
| Superfrost Plus slides | Thermo-Scientific® | |
| **Software and algorithms** | | |
| QuPath v0.4.3 software | https://qupath.github.io/ | |
| R Studio, with R 4.4.2 | https://www.r-project.org/ | |
| R-package Rcmdr | https://cran.r-project.org/web/packages/Rcmdr/index.html | |
| GraphPad Prism software 9 | https://www.graphpad.com/ | |
